# Comparative analyses of SARS-CoV-2 binding (IgG, IgM, IgA) and neutralizing antibodies from human serum samples

**DOI:** 10.1101/2020.08.10.243717

**Authors:** Livia Mazzini, Donata Martinuzzi, Inesa Hyseni, Giulia Lapini, Linda Benincasa, Pietro Piu, Claudia Maria Trombetta, Serena Marchi, Ilaria Razzano, Alessandro Manenti, Emanuele Montomoli

**Author notes:** Corresponding authors: Alessandro Manenti, Vismederi Resrarch S.r.l. 53100 Siena, Italy;, Livia Mazzini, VisMederi S.r.l. 53100 Siena, Italy.

## Abstract

A newly identified coronavirus, named SARS-CoV-2, emerged in December 2019 in Hubei Province, China, and quickly spread throughout the world; so far, it has caused more than 18 million cases of disease and 700,000 deaths. The diagnosis of SARS-CoV-2 infection is currently based on the detection of viral RNA in nasopharyngeal swabs by means of molecular-based assays, such as real-time RT-PCR. Furthermore, serological assays aimed at detecting different classes of antibodies constitute the best surveillance strategy for gathering information on the humoral immune response to infection and the spread of the virus through the population, in order to evaluate the immunogenicity of novel future vaccines and medicines for the treatment and prevention of COVID-19 disease. The aim of this study was to determine SARS-CoV-2-specific antibodies in human serum samples by means of different commercial and in-house ELISA kits, in order to evaluate and compare their results first with one another and then with those yielded by functional assays using wild-type virus. It is important to know the level of SARS-CoV-2-specific IgM, IgG and IgA antibodies in order to predict population immunity and possible cross-reactivity with other coronaviruses and to identify potentially infectious subjects. In addition, in a small sub-group of samples, we performed a subtyping Immunoglobulin G ELISA. Our data showed an excellent statistical correlation between the neutralization titer and the IgG, IgM and IgA ELISA response against the receptor-binding domain of the spike protein, confirming that antibodies against this portion of the virus spike protein are highly neutralizing and that the ELISA Receptor-Binding Domain-based assay can be used as a valid surrogate for the neutralization assay in laboratories which do not have Biosecurity level-3 facilities.

## INTRODUCTION

Coronaviruses (CoVs) are enveloped, positive single-stranded RNA viruses belonging to the *Coronaviridae* family. Several members of this family, such as human coronavirus (HCoV) OC43, NL63 and 229E, cause mild common colds every year in the human population [1]. Three highly pathogenic novel CoVs have appeared in the last 18 years; Severe Acute Respiratory Syndrome (SARS-CoV-1) emerged in November 2002 in Guangdong province, causing more than 8,000 confirmed cases and 774 deaths [2], [3], Middle East Respiratory Syndrome (MERS-CoV) was discovered in June 2012 [4], and Severe Acute Respiratory Syndrome Coronavirus 2 (SARS-CoV-2) emerged in Wuhan, Hubei province, China, in December 2019; this last was declared a pandemic on March 11^th^ 2020 by the World Health Organization (WHO). The global impact of the SARS-CoV-2 outbreak, with almost 18 million cases and 700,000 deaths reported (as of August 7^th^ 2020) [5], is unprecedented. Several data have confirmed that the infection initially arose from contact with animals in the Hunan seafood market. Subsequently, human-to-human transmission occurred, leading to a very high rate of laboratory-confirmed infection in China [6], [7]. Precise diagnosis of Coronavirus disease (COVID-19) is essential in order to promptly identify infected individuals, to limit the spread of the virus and to allow those who have been infected to be treated in the early phases of the infection. To date, real-time polymerase chain reaction (RT-PCR) is the most widely employed method of diagnosing COVID-19. However, rapid, large-scale testing has been prevented by the high volume of demand and the shortage of the materials needed for mucosal sampling [8]. Standardized serological assays able to measure in a specific manner the antibody responses may help to overcome these problems and may support a significant number of relevant applications. Indeed, serological assays are the basis on which to establish the rate of infection (severe, mild and asymptomatic) in a given area, to calculate the percentage of the population susceptible to the virus and to determine the fatality rate of the disease. It has been demonstrated in a non-human primate model [9] that, once the antibody response has been established, re-infection and, consequently, viral shedding, is unlikely. Furthermore, serological assays can help to identify subjects with strong antibody responses, who could serve as donors for the generation of monoclonal antibody therapeutics [10]; in addition, these assays can be used to define future correlates of protection for SARS-CoV-2. The spike glycoprotein (S-protein), a large transmembrane homotrimer of approximately 140kDa, has a pivotal role in viral pathogenesis, mediating binding to target cells through the interaction between its receptor-binding domain (RBD) [11] and the human angiotensin converting enzyme 2 (ACE2) receptor. The S-protein has been found to be highly immunogenic, and the RBD is considered the main target in the effort to elicit potent neutralizing antibodies [12], [13]. Two subunits give rise to the S-protein: S1, which mediates attachment, and S2, which mediates membrane fusion. The CoV S-protein is a class I fusion protein, and protease cleavage is required for activation of the fusion process [14].

Few data documenting the differences in systemic immunoglobulin G (IgG) and their subclasses, IgM and IgA, in terms of their responses against SARS-CoV-2, are as yet available, especially data comparing these responses with the neutralizing one. It is well recognized that IgG levels are crucial to protection from viral disease [15]. In humans, the four subclasses of IgG (IgG1, IgG2, IgG3, IgG4) differ in function [16]. Importantly, IgG1 and IgG3 play a key role in many fundamental immunological functions, including virus neutralization, opsonization and complement fixation [17].

We conducted this comparative study for two purposes: first, to investigate the sensitivity and specificity of detection of different ELISA kits compared with Micro-Neutralization (MN) results; second, to investigate the difference in spike-RBD-specific IgG, IgM and IgA antibody responses in human serum samples.

## RESULTS

### Set-up and standardization of in-house ELISAs

Several purified recombinant S-proteins (S1 and RBD domain) were tested for their ability to detect specific human antibodies: S1-SARS-CoV-2 (HEK293) (from Native Antigen); S1-SARS-CoV-2 (HEK293) (from eEnzyme); S1-SARS-CoV-2 (HEK293) (from ACROBiosystems); Spike RBD-SARS-CoV-2 (Baculovirus-Insect and HEK293) (from Sino Biological). Each protein was evaluated on using three coating concentrations (1, 2 and 3 µg/mL) and four different dilutions of the secondary Horse Radish Peroxidase (HRP) conjugate anti-human IgG, IgM and IgA antibodies. We also evaluated the impact of the incubation time of the HRP by incubating the plates for 1 hour or 30 minutes, and concluded that the best and clearest signal was always seen after the shortest incubation. To set the assays, three human serum samples derived from convalescent donors, along with a pool of Micro Neutralization Titres (MNT) and ELISA (commercial Kit)-negative human serum samples, were used as positive and negative controls, respectively. As a test control, human IgG1 monoclonal antibody (mAb) anti-SARS-CoV-2 spike (S1) (CR3022 Native antigen), human IgM mAb anti-SARS-CoV-2 spike (S1) (CR3022 Absolute antibody) and human IgG1 anti-Spike RBD (SCV2-RBD eEnzyme) were used. Additionally, several human sera hyper-immune to various infectious diseases (influenza, diphtheria and pertussis) were used to assess the specificity of the assay in detecting only antibodies against SARS-CoV-2 S1 or the RBD protein. Alternative blocking/diluent solutions containing 1% Bovine Serum Albumin (BSA), 2.5% milk and 5% milk were tested. The specificity of the test increased significantly on using the 5% milk blocking solution in comparison with BSA, which occasionally yielded non-specific results and displayed a generally higher background. Finally, the two proteins that yielded the best results in terms of sensitivity and specificity were chosen as candidates for the tests: the purified S1-protein (HEK derived) from eEnzyme and the Purified RBD protein (HEK derived) from Sino Biological.

### Correlation between ELISAs and Neutralization

Each serum sample was tested by means of *in-house* ELISA S1 and RBD-specific IgG, IgM and IgA (VM_IgG_S1, VM_IgG_RBD, VM_IgM_S1, VM_IgM_RBD, VM_IgA_RBD) and by means of the Euroimmun S1 Commercial ELISA kit, along with the functional MNT assay. The distribution of the MNT was strongly asymmetric, with most of the values (153/181) being equal to 5 (i.e. negative). The other values observed (from 10 to 1280 in a 2-fold dilution series) were uniformly distributed. Concerning the ELISA S1, we performed two different tests: one by means of a commercial (Euroimmun) kit and the other an in-house ELISA. According to Spearman’s rank correlation coefficients and statistical significance (Tables 1 and 2), we registered the highest agreement between the ELISA VM_IgG_RBD and MNT, and between the VM_IgA_RBD and MNT, with coefficients of 0.83 and 0.85, respectively. The lowest correlations were found for ELISA Euroimmun vs MNT, and for VM_IgG_S1 vs MNT, with coefficients of 0.49 and 0.45, respectively. As can be seen from the correlation plot (Figure 1), the IgA response was closely linked with a positive MNT response. Moreover, on dissecting all the results for each serum sample (data not shown), we noted that, in those subjects in whom we registered a high neutralization titer, we always observed a positive IgA signal.

**Table 1.** Spearman’s rank correlation coefficients

|  | *MNT* | *EUROIMMUN* | *VM_IgG_S1* | *VM_IgG_RBD* | *VM_IgM_S1* | *VM_IgM_RBD* | *VM_IgA_RBD* |
| --- | --- | --- | --- | --- | --- | --- | --- |
| *MNT* | 1.00 | 0.49 | 0.45 | 0.83 | 0.52 | 0.73 | 0.85 |
| *EUROIMMUN* | 0.49 | 1.00 | 0.77 | 0.59 | 0.50 | 0.54 | 0.51 |
| *VM_IgG_S1* | 0.45 | 0.77 | 1.00 | 0.59 | 0.43 | 0.44 | 0.53 |
| *VM_IgG_RBD* | 0.83 | 0.59 | 0.59 | 1.00 | 0.49 | 0.73 | 0.84 |
| *VM_IgM_S1* | 0.52 | 0.50 | 0.43 | 0.49 | 1.00 | 0.50 | 0.45 |
| *VM_IgM_RBD* | 0.73 | 0.54 | 0.44 | 0.73 | 0.50 | 1.00 | 0.69 |
| *VM_IgA_RBD* | 0.85 | 0.51 | 0.53 | 0.84 | 0.45 | 0.69 | 1.00 |

**Table 2.** Statistical significance of Spearman’s rank correlation coefficients

|  | *MNT* | *EUROIMMUN* | *VM_IgG_S1* | *VM_IgG_RBD* | *VM_IgM_S1* | *VM_IgM_RBD* | *VM_IgA_RBD* |
| --- | --- | --- | --- | --- | --- | --- | --- |
| *MNT* | 0.0E+00 | 3.8E-07 | 3.3E-10 | 8.1E-47 | 3.5E-14 | 1.5E-31 | 7.0E-53 |
| *EUROIMMUN* | 3.8E-07 | 0.0E+00 | 2.8E-20 | 1.9E-10 | 1.8E-07 | 1.5E-08 | 1.3E-07 |
| *VM_IgG_S1* | 3.3E-10 | 2.8E-20 | 0.0E+00 | 3.4E-18 | 2.1E-09 | 5.8E-10 | 2.1E-14 |
| *VM_IgG_RBD* | 8.1E-47 | 1.9E-10 | 3.4E-18 | 0.0E+00 | 1.5E-12 | 3.1E-31 | 2.5E-50 |
| *VM_IgM_S1* | 3.5E-14 | 1.8E-07 | 2.1E-09 | 1.5E-12 | 0.0E+00 | 5.5E-13 | 3.1E-10 |
| *VM_IgM_RBD* | 1.5E-31 | 1.5E-08 | 5.8E-10 | 3.1E-31 | 5.5E-13 | 0.0E+00 | 2.9E-27 |
| *VM_IgA_RBD* | 7.0E-53 | 1.3E-07 | 2.1E-14 | 2.5E-50 | 3.1E-10 | 2.9E-27 | 0.0E+00 |

**Figure 1.**
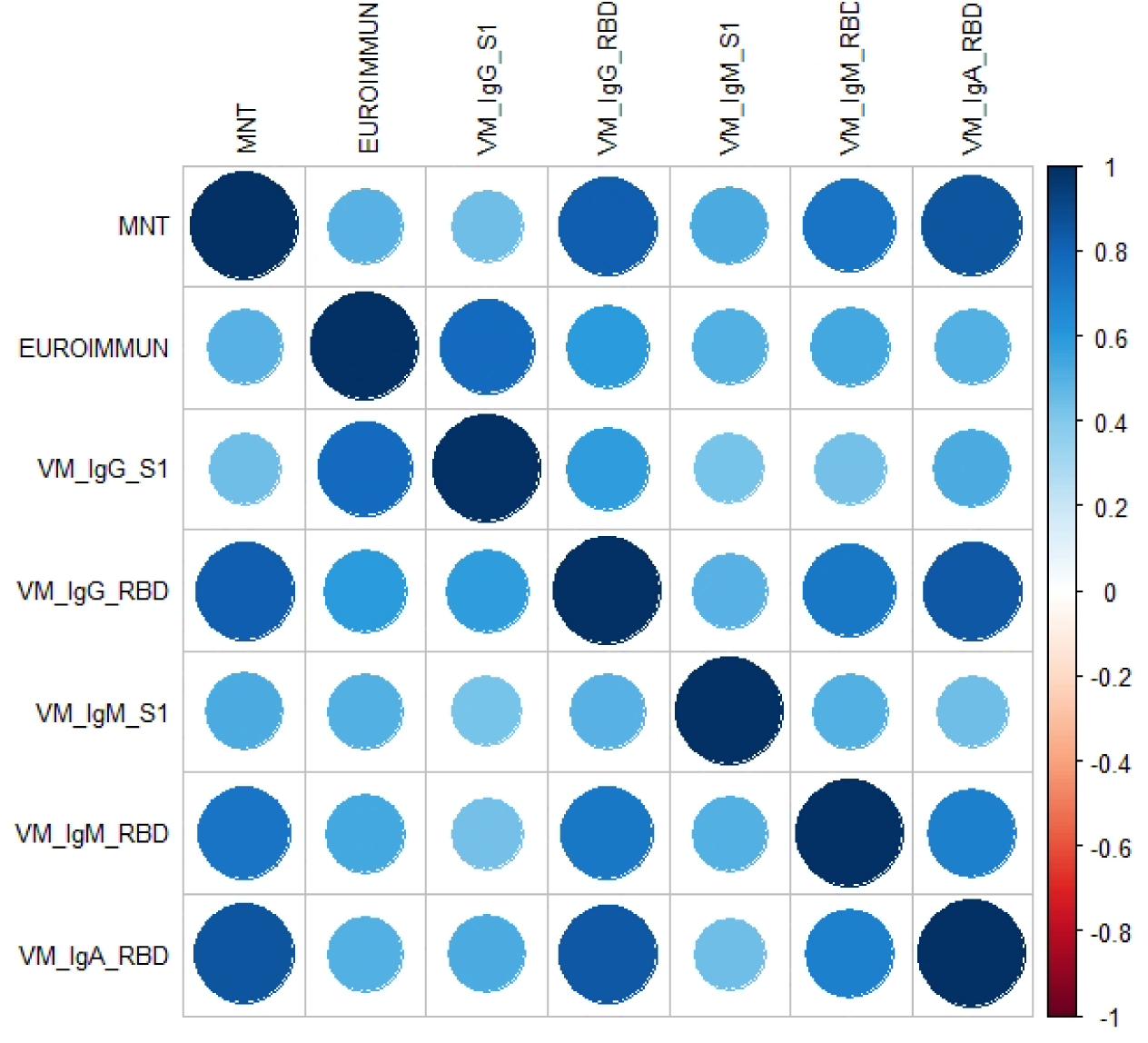
The correlation plot associated to the measured coefficients of Spearman’s rank correlation. The magnitude of the coefficients is represented by circles and a color gradient: the larger the area of the circle and the more intense the tone of the color, the greater the correlation. The direction of the correlation is indicated by the color scale: blue tones for positive correlations and red tones for negative correlations.

Interestingly, in 9 MNT-positive samples, we found a complete absence of S1 signal on using Euroimmun, VM_IgG_S1 and VM_IgM_S1 ELISA kits but, on the other hand, high and detectable IgG and IgM RBD-specific signals.

### Classification analysis: Elastic Net

Over a training set of data, the optimal hyper-parameters estimated for the Elastic Net (EN) model were lambda=0.0136 and alpha=0.76, which minimized the error of cross-validation (= 0.3809). The EN model selected three significant predictors of the MNT results, namely VM-IgG-RBD, VM-IgM-RBD, and VM-IgA-RBD; the estimates of their coefficients were 0.0035, 0.0060 and 0.0013, respectively, while the intercept of the model was -2.9741. These results were entered into the score function, whereby we predicted the MN titers. From the ROC curve (Figure 2A), we evaluated the performance of the predictions in terms of sensitivity and specificity. On balancing sensitivity and specificity, we obtained the optimal cut-off of 0.092, with sensitivity = 85.7 % (95% CI = [42.1% – 99.6%]) and specificity = 98.1% (95% CI = [89.9% – 99.6%]) (Figure 2B). Overall, these findings indicated that the Vismederi RBD-based ELISA methods were highly accurate and, particularly, presented the features of a highly specific diagnostic test when jointly considered.

**Figure 2.**
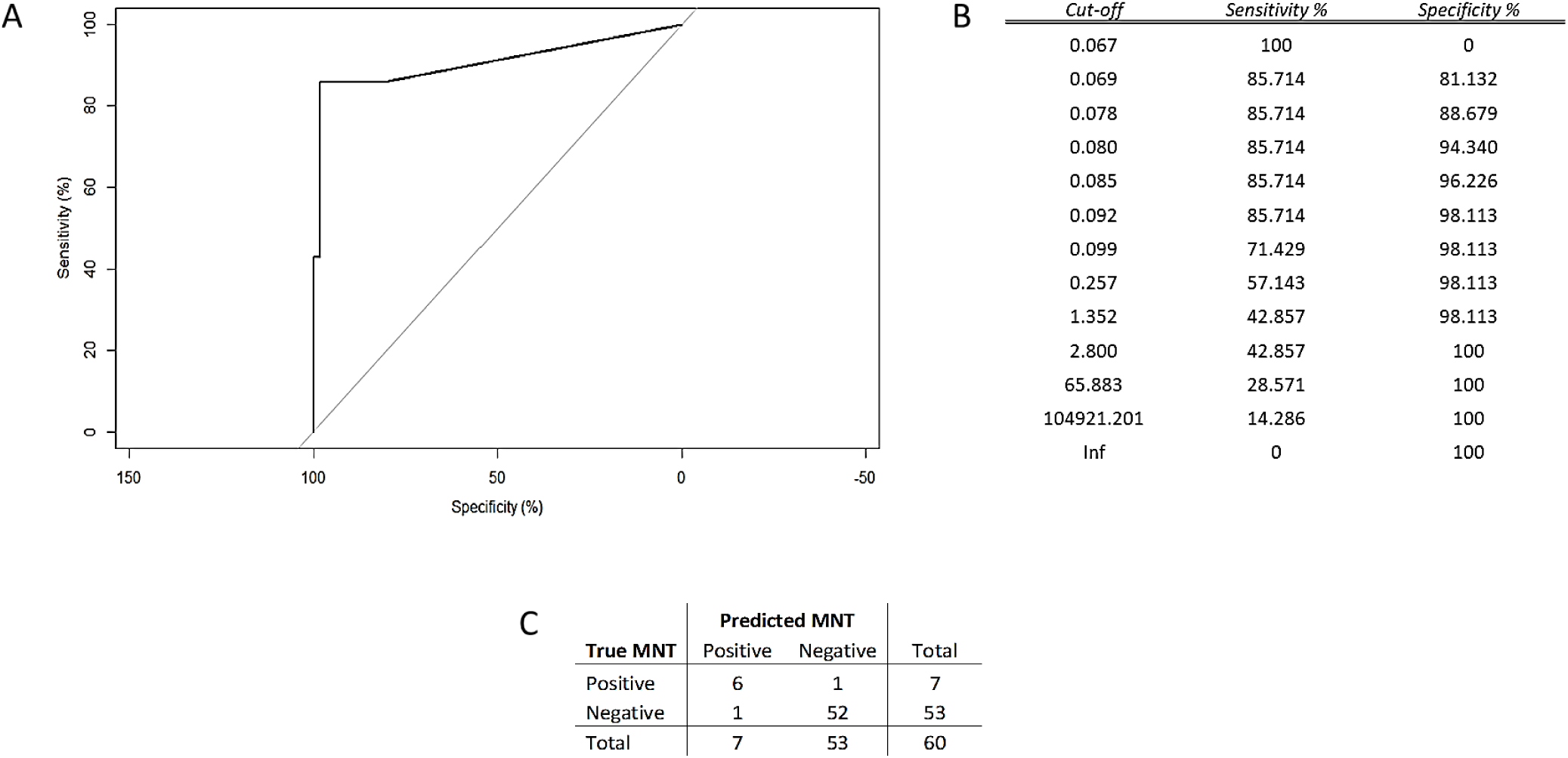
A) Analysis of the ROC curve referred to the test set proved that the results of the EN model attained high accuracy in predicting the MNT values. Measurement of the area under the curve, AUC = 90.7 %, supported this conclusion; B) Summary table of ROC analysis; C) Error matrix.

The samples which yielded a score below the cut-off identified were classified as “negative”, and the remaining samples as “positive”. We then compared these predictions with the known results of the test-set (Figure 2C).

Analysis of the error matrix indicated an overall Accuracy (ACC) of 96.7% (95% CI = [88.5% – 99.6%]), and a No Information Rate (NIR) of 88.3% (95% CI = [77.4% – 95.2%]). Since the ACC was significantly higher than the NIR (p = 0.02), we may claim that the model built with the In-house (VM) RBD-based ELISAs conveyed effective information. The extremely high value of the odds ratio (OR) = 312.0, (95% CI = [17.2 – 5657.7]) revealed the strong association between the MNT results and the model predictions. Specifically, the positive predictions were 312 times more likely to occur in association with positive MNT than the negative predictions.

### IgG subtyping of serum samples

We also evaluated the ELISA IgG subtyping response (IgG1, IgG2, IgG3, IgG4) in a small subgroup (14) of MN-positive samples. ELISA plates were coated with RBD purified antigen. Our results, although derived from a small group of subjects, are in line with previous findings by Amanat and colleagues [18]. Strong reactivity for IgG1 and IgG3 was found in almost all samples, with the IgG3 subclass showing the highest percentage of detection. Low and very low reactivity was found for IgG4 and IgG3, respectively (Figure 3).

**Figure 3:**
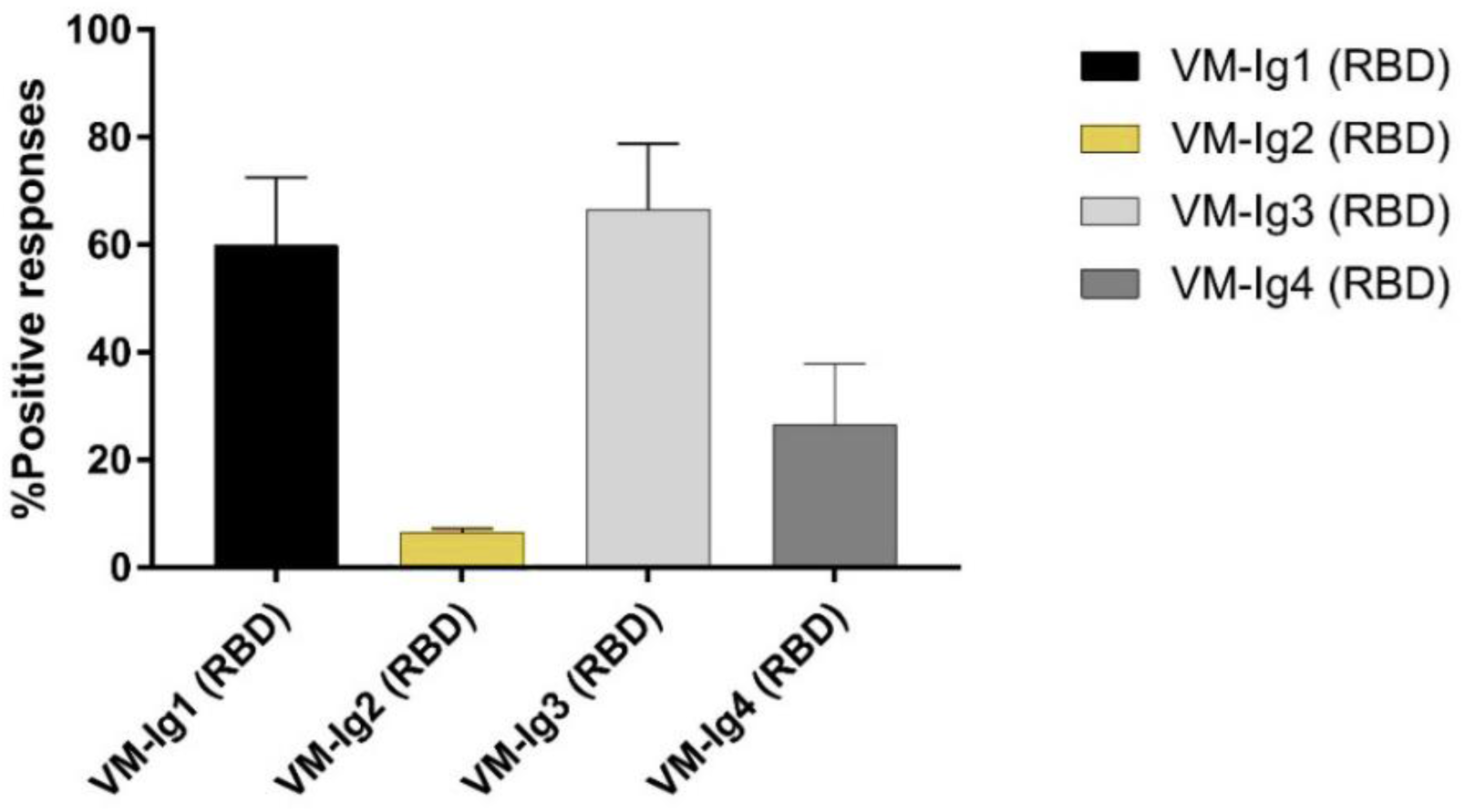
Percentage of detection of IgG1, IgG2, IgG3 and IgG4 in all 14 human samples positive on Micro-neutralization assay.

## DISCUSSION

Like most of the emerging infectious diseases that affect humans, this new HCoV also originated from animals [19], [20]. Owing to the rapid increase in some human practices, such as deforestation, urbanization and the husbandry of wild animal species, over the years the emergence of new pathogens has become an extremely serious problem. The rapid global spread of the novel SARS-CoV-2 is posing a serious health threat to the entire world. There is now an urgent need for well-standardized serological assays that can detect different classes of antibodies against the novel coronavirus, and which can be used alongside the classical diagnostic molecular methods (RT-PCT). Indeed, owing to the huge demand in recent months, the availability of the reagents and equipment needed to promptly carry out analyses is still inadequate. Moreover, if sample collection and storage are improperly conducted, molecular tests may yield false-negative results in subjects who carry the virus [21]. Previous studies on SARS-CoV-1 have shown that virus-specific IgG (and IgM) levels can be a valid surrogate for serological diagnosis [22], [23]. Indeed, the present study had two major goals: a) to standardize and make as reliable as possible ELISAs able to detect different classes of immunoglobulins, and b) to broaden the data-set of information on comparisons between the results of different serological tests, which could be precious for future evaluations of serological diagnoses and vaccine assessments [24]. Specifically, in this study, ELISA results were always compared with those of the functional assay (MN), which is commonly assumed as a benchmark and the gold standard. Since its first isolation and characterization, this new HCoV strain has been classified, according to the WHO guidelines, as a biosafety level 3 (BSL3) pathogen. This has placed some limits on the implementation of neutralization tests, as relatively few laboratories have level-3 biocontainment facilities. The ELISAs are a good surrogate for the MN assay in terms of sensitivity, safety and throughput [25]–[27]. However, it is very important to evaluate and estimate the best antigen/s to use in these platforms in order to obtain a reliable and similar response to that of the neutralization test, which indicates the functional response. This is why we compared all our results with those of the MN. As in the case of influenza hemagglutinin [28], antibodies specific to the RBD domain of the S-protein are the key to viral neutralization. In this study, together with the IgG, IgM and IgA analyses, we also evaluated the responses of IgG subclasses in those subjects who showed both a high RBD ELISA signal and proven neutralization activity. Our results are in line with previous findings [18] and confirm IgG1 and IgG3 as the subtypes with the strongest reactivity in all samples [29]. Only in a small number of subjects did we find IgG2 and IgG4 responses. IgG1 and IgG3 are involved in critical immunologic functions, such as neutralization, opsonization, complement fixation and antibody-dependent cellular cytotoxicity (ADCC). On the other hand, IgG2 plays an important role in protecting against infection by encapsulated microorganisms [30]; IgG4 is generally a minor component of the total immunoglobulin response and is induced in response to continuous antigenic stimulation [31, p. 4].

Regarding the ELISA IgG, IgM and IgA, the main results can be summarized as follows: a) all the proposed statistical analyses indicated a close relationship between the results of MNT and in-house RBD-based ELISAs, namely VM-IgG-RBD, VM-IgM-RBD and VM-IgA-RBD; these results are in line with previous reports by Amanat and colleagues [18], [32]; b) the cross-validation technique applied to the EN model allowed us to obtain robust results. In the out-of-sample data (i.e., the randomly chosen test-data) highly accurate, and, particularly, highly specific performance was observed; c) in large-scale screening operations, it is very important to have a highly specific test, as this guards against the risk of misclassification of true-negative samples with a wide margin of certainty. A highly specific test is particularly useful in order to confirm a diagnosis already made by means of other methods, and when a false-positive result would have a great impact. Indeed, a highly specific test is of most help to the clinician when it provides a positive result.

An overview of all the results yielded by ELISA and MNT (data not shown) reveals that the highest neutralization activity against SARS-CoV-2 is achieved when all three immunoglobulins, IgG, IgM and IgA are detected, as if to indicate the presence of a synergistic or additive effect among different classes of antibodies. This observation can be explained by the fact that the human population is completely naïve for SARS-CoV-2 and that IgG or IgM alone is not able to mount an ideal neutralizing immune response. Indeed, one of the most important features of adaptive immunity is the generation of immunological memory and the ability of the immune system to learn from its experiences of encounters with the same pathogen, thereby becoming more effective over time [33].

Interestingly, in nine samples, neither in-house nor commercial kits detected any IgG and IgM signal for the S1 protein, while a strong signal for RBD-specific IgG, IgM and IgA was detected. As all nine samples displayed exactly same trend, it seems that these results could be due to the folding of the three-dimensional S1 protein structure after the production in HEK293 cells, which could have masked some epitopes recognized by the antibodies expressed in these nine subjects. By contrast, these epitopes may be well exposed in the RBD protein and can be bound by antibodies, which would explain the differences in signals.

To conclude, these results confirm what has already been reported [34], i.e. that the immune response to SARS-CoV-2 is very variable, but that antibodies targeting the RBD domain have the highest probability of being neutralizing, and in some cases of being strongly neutralizing. The present study constitutes preliminary research into the development of an ELISA that can semi-quantify anti-SARS-CoV-2 human antibodies in a specific and repeatable way. The next step will be to completely validate these ELISAs according to the criteria established by the International Council for Harmonization of Technical Requirements for Pharmaceuticals for Human Use [35, p. 2], and to analyze the performance and specificity of these tests with specific human serum samples that are highly positive towards different HCoVs.

## MATERIALS AND METHODS

### Serum samples

In March/April 2020, 181 human serum samples were collected by the laboratory of Molecular Epidemiology of the University of Siena, Italy. The samples were anonymously collected in compliance with Italian ethics law.

Three human serum samples from confirmed cases of COVID-19 were kindly provided by Prof. Valentina Bollati from the University of Milan. Human IgG1 anti-SARS-CoV-2 Spike (S1) Antibody CR3022 (Native Antigen, place), Human IgM anti-SARS-CoV-2 Spike (S1) Antibody CR3022 (Native Antigen, Oxford, UK) and anti-Spike RBD (SARS-CoV-2/COVID 19) human monoclonal antibody (eEnzyme, Gaithersburg, USA) were used as positive controls in ELISA. Human serum minus (IgA/IgM/IgG) (Cod. S5393, Sigma, St. Louis, USA) was also used as a negative control in MNT and ELISA.

Three human serum samples containing heterologous neutralizing antibodies, provided by NIBSC, were used to verify the specificity of the ELISA: WHO 1^st^ International Standard for Pertussis antiserum (lot. 06/140); WHO 2^nd^ International Standard for antibody to influenza H1N1pdm virus (lot. 10/202); WHO 1^st^ International Standard for Diphtheria Antitoxin (lot: 10/262).

### Cell Culture

Vero E6 cells, acquired from the American Type Culture Collection (ATCC - CRL 1586), were cultured in Dulbecco’s Modified Eagle’s Medium (DMEM) - High Glucose (Euroclone, Pero, Italy) supplemented with 2 mM L-Glutamine (Lonza, Milan, Italy), 100 units/mL penicillin-streptomycin mixture (Lonza, Milan, Italy) and 10% of Fetal Bovine Serum (FBS), at 37°C, in a 5% CO_2_ humidified incubator.

VERO E6 cells were seeded in a 96-well plate using D-MEM high glucose 2% FBS at a density of 1.5 x 10^6^cells per well, in order to obtain a 70-80% sub-confluent cell monolayer after 24 hours.

### SARS-CoV-2 purified antigen, live virus and titration

Five different purified recombinant S proteins (S1 and RBD domain) were tested for their ability to detect specific human antibodies: S1-SARS-CoV-2 (HEK293) Cod. REC31806-500, (Native Antigen, Oxford, UK); S1-SARS-CoV-2 (HEK293) Cod. SCV2-S1-150P (eEnzyme Gaithersburg, MD, USA); S1-SARS-CoV-2 (HEK293) Cod. S1N-C52H3 (ACROBiosystems, Newark, DE, USA); Spike RBD-SARS-CoV-2 (Baculovirus-Insect cells) Cod. 40592-V08B and (HEK293) Cod. 40592-V08H (Sino Biological, Beijing, China).

SARS CoV-2 - strain *2019-nCov/Italy-INMI1 –* wild-type virus was purchased from the European Virus Archive Global (EVAg, Spallanzani Institute, Rome). The virus was titrated in serial 1-log dilutions to obtain a 50% tissue culture infective dose (TCID50) on 96-well culture plates of VERO E6 cells. The plates were observed daily for the presence of cytopathic effect (CPE) by means of an inverted optical microscope for a total of 4 days. The end-point titers were calculated according to the Spearman-Karber formula [36].

### Micro-neutralization assay

The MNT was performed as previously reported by Manenti et al. [37]. Briefly, 2-fold serial dilutions of heat-inactivated serum samples were mixed with an equal volume of viral solution containing 100 TCID50 of SARS-CoV-2. The serum-virus mixture was incubated for 1 hour at 37°C in a humidified atmosphere with 5% CO_2_. After incubation, 100 µl of the mixture at each dilution was passed to a 96-well cell plate containing a 70-80% confluent VERO E6 monolayer. The plates were incubated for 3 days at 37°C in a humidified atmosphere with 5% CO_2_. After the incubation time, each well was inspected by means of an inverted optical microscope to evaluate the percentage of CPE. The highest serum dilution that protected more than 50% of cells from CPE was taken as the neutralization titer.

### Commercial Enzyme-Linked Immunosorbent Assay (ELISA)

Specific anti-SARS-CoV-2 IgG antibodies were detected by means of two commercial ELISA kits: Euroimmun and Eagle Biosciences.

Euroimmun-ELISA plates were coated with recombinant structural protein (S1 domain) of SARS-CoV-2. The assay provides semi-quantitative results by calculating the ratio of the optical density (OD) of the serum sample over the OD of the calibrator. According to the manufacturer’s instructions, positive samples have a ratio ≥ 1.1, borderline samples a ratio between 0.8 and 1.1 and negative samples a ratio < 0.8.

### In-House S1 and RBD Enzyme-Linked Immunosorbent Assay (ELISA) IgG, IgM and IgA

ELISA plates were coated with 1µg/mL of purified recombinant Spike S1 Protein (aa 18-676) (eEnzyme, Gaithersburg, MD, USA) or with 1µg/mL Spike-RBD (Arg319-Phe541) (Sino Biological, China), both expressed and purified from HEK 293 cells. After overnight incubation at +4°C, coated plates were washed three times with 300 µl/well of ELISA washing solution containing Tris Buffered Saline (TBS)-0.05% Tween 20, then blocked for 1 hour at 37°C with a solution of TBS containing 5% of Non-Fat Dry Milk (NFDM; Euroclone, Pero, Italy).

Serum samples were heat-inactivated at 56°C for 1 hour in order to reduce the risk of the presence of live virus in the sample. Subsequently, 3-fold serial dilutions, starting from 1:100 in TBS-0.05% Tween 20 5% NFDM, were performed up to 1:2700. Plates were washed three times, as previously; then 100 µl of each serial dilution was added to the coated plates by means of a multichannel pipette and incubated for 1 hour at 37°C. Next, after the washing step, 100 µl/well of Goat anti-Human IgG-Fc HRP-conjugated antibody or IgM (µ-chain) and IgA (α-chain) diluted 1:100,000 or 1:100,000 and 1:75,000, respectively, (Bethyl Laboratories, Montgomery USA) were added. Plates were incubated at 37°C for 30 minutes. Following incubation, plates were washed and 100 µl/well of 3,3′,5,5′-Tetramethylbenzidine (TMB) substrate (Bethyl Laboratories, Montgomery, USA) was added and incubated in the dark at room temperature for 20 minutes. The reaction was stopped by adding 100 µl of ELISA stop solution (Bethyl Laboratories, Montgomery, USA) and read within 20 minutes at 450 nm. To evaluate the OD a SpectraMax ELISA plate reader was used. A cut-off value was established as 3 times the average of OD values from blank wells (background: no addition of analyte).

Samples with ODs below the cut-off value at the first (1:100) dilution were assigned as negative, samples with titres between 100 and 300 were assigned as positive, and samples showing a titer above 300 were assigned as highly positive.

### In-House RBD Enzyme-Linked Immunosorbent Assay (ELISA) IgG1, IgG2, IgG3 and IgG4

An indirect ELISA was performed in order to determine the RBD-specific IgG1, IgG2, IgG3 and IgG4 antibody concentration in serum samples [38]. 96-well plates were coated with 1µg/mL of purified Spike-RBD (Sino Biologicals). Serum samples were diluted from 1:50 to 1:400. Mouse anti-human IgG1, IgG2, IgG3 and IgG4 Fc-HRP (Southern Biotech, USA) secondary antibodies were used at 1:8000 dilution. The cut-off values were established as reported above.

### Statistical analysis

Spearman’s rank correlation analysis enabled us to determine whether, and to what extent, the MNT assay was associated with the ELISAs. A classification analysis gave further insight into the relationship between the MNT results and the in-house ELISAs. We defined the MN as the target variable and recoded its results by assigning the label “0” to values of 5, and the label “1” otherwise. We implemented an elastic net (EN) to classify the MNT results. The EN is a rather sophisticated generalized linear model (GLM), which addresses the issues caused by multi-collinearity among predictors. We set the binomial family for the GLM after dichotomizing the variable MNT; therefore, we followed a logistic-like model approach in the implementation of the EN. The EN produces a selection of the variables based on a convex penalty function, which is a combination of the ridge regression and the LASSO (Least Absolute Shrinkage and Selection Operator) penalties, say l1 and l2 respectively, controlled by the hyper-parameter alpha = l2/(l1+l2). The hyper-parameter, lambda, by contrast, regulates the level of penalization in the model [39]. To improve the generalization capability of the EN, we trained the model over a randomly selected subset of data (121/181) and verified its robustness over an independent subset of the residual data (60/181), which did not enter the model during the training stage. The cross-validation technique prevented the occurrence of over-fitting problems in the estimates. On the base of the values of the predictors of the test set, X, and their estimated EN coefficients, b, we built a score function, S, as follows:

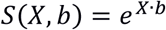

The probability of a positive MNT assignment for the predicted results was then expressed as:

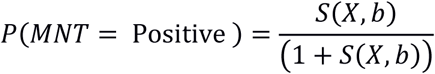

We calculated the performance of the EN in terms of sensitivity, i.e., the percentage of positive MNT correctly predicted, and specificity, i.e., the percentage of negative MNT correctly predicted, and represented their related Receiver Operating Characteristic (ROC) curve. The optimal combination of sensitivity and specificity enabled us to detect the cut-off in the score function; test samples were classified as positive if their score was above this cut-off value and as negative if the score was below it, with the minimum error probability.

## ACKNOWLEDGMENTS

We thank the Laboratory of Molecular and Developmental Medicine of the University of Siena for providing the human serum samples. Furthermore, we thank Dr. Valentina Bollati from the University of Milan for providing the serum samples from COVID-19-positive patients. This publication was supported by the European Virus Archive Global (EVAg) project, which has received funding from the European Union’s Horizon 2020 research and innovation program under grant agreement No 653316.

## Authors’ Contributions

LM performed the set-up experiments and standardized *in-house* ELISAs; LM, DM and HY performed all the ELISA experiments; PP evaluated the results and performed the statistical analyses; LB and IR handled the Vero E6 cells and prepared the plates for neutralization experiments; AM performed the Micro-neutralization experiments; CMT and SM performed the Euroimmun assay at the University site and provided the human serum samples; AM and GL designed the experiments; AM and LM prepared the draft of the manuscript; EM supervised the study. All authors have approved the final version of the manuscript.

